# Towards harmonized spectral quantification in MRSI: Comparative analysis of Backward-Linear-Predicted and original ¹H-FID-MRSI dephased data

**DOI:** 10.64898/2025.12.05.692527

**Authors:** Alessio Siviglia, Brayan Alves, Cristina Cudalbu, Bernard Lanz

## Abstract

**Object:** We hypothesized that the inherent acquisition delay (AD) in ¹H-FID-MRSI can introduce systematic LCModel quantification biases due to strong spectral dephasing, and that Backward-Linear-Prediction (BLP) reconstruction toward AD = 0 ms can harmonize metabolite estimates across acquisitions with various delays.

**Materials and Methods:** 2D ¹H-FID-MRSI were acquired in rats at 14.1T with three AD values (0.71, 0.94, 1.30 ms). Hippocampal metabolites were quantified using LCModel and AD-matched basis sets. Complementary Monte-Carlo simulations (n = 1000) replicated ¹H-FID-MRSI spectra at multiple ADs under realistic SNR conditions. BLP was applied to *in vivo* and simulated FIDs to back-predict missing points up to AD = 0 ms, enabling quantification within a unified basis set framework.

**Results:** *In vivo* and simulated data showed clear AD-dependent variations for several metabolites (Gln, tCho, tNAA, Ins, Tau), with discrepancies frequently >10% despite AD-specific basis sets. Simulations confirmed metabolite-specific biases increasing with AD. BLP reconstruction preserved quantification consistency up to ∼0.98 ms of recovered FIDs, reducing inter-AD mismatches *in vivo*—particularly for Tau, tNAA and tCho—lowering the mean discrepancy from 10.5% to ∼5%.

**Discussion:** These findings show that AD affects ¹H-FID-MRSI quantification in LCModel, whereas BLP reconstruction can harmonize spectra across delays by enabling a virtual AD = 0 ms quantification scheme. This supports BLP as a practical strategy to improve consistency and comparability in MRSI studies.

## 1. Introduction

Proton free-induction-decay magnetic resonance spectroscopic imaging (^1^H-FID-MRSI) has shown great potential for neurological studies. The MRSI technique enables the characterization of localized magnetic resonance spectra of brain metabolites non-invasively. This provides *in vivo* mapping of metabolites concentration over the brain, establishing itself over the last decade as a powerful methodology for human [1] [2] [3] and preclinical research [4]. ^1^H-FID-MRSI offers significant technical advantages such as minimal signal loss due to T2 relaxation and J-evolution, as well as no in-plane chemical shift displacement errors [4] [5] [6]. At ultra-high field, good levels of SNR and a wide set of detectable metabolites are provided by a relatively simple sequence design, which can further offer short acquisition times when working with short repetition times (TR) inpartially T_1_-saturated acquisitions [5] [7] [8].

However, a limitation of ^1^H-FID-MRSI lies in the inherent delay between the excitation pulse and the start of the FID acquisition, which is needed to spatially encode the signal (Fig. 1a). This acquisition delay (AD, defined as the sum of the excitation pulse length fraction, slice-rephasing gradient and phase-encoding duration (applied concomitantly), and the dead time (ADC initialisation)), varies depending on the system performance and the protocol, typically ranging around ∼ 1 ms [1] [3] [4] [9]. This delay results in a strong first order phase in the acquired spectrum.

**Fig. 1:**
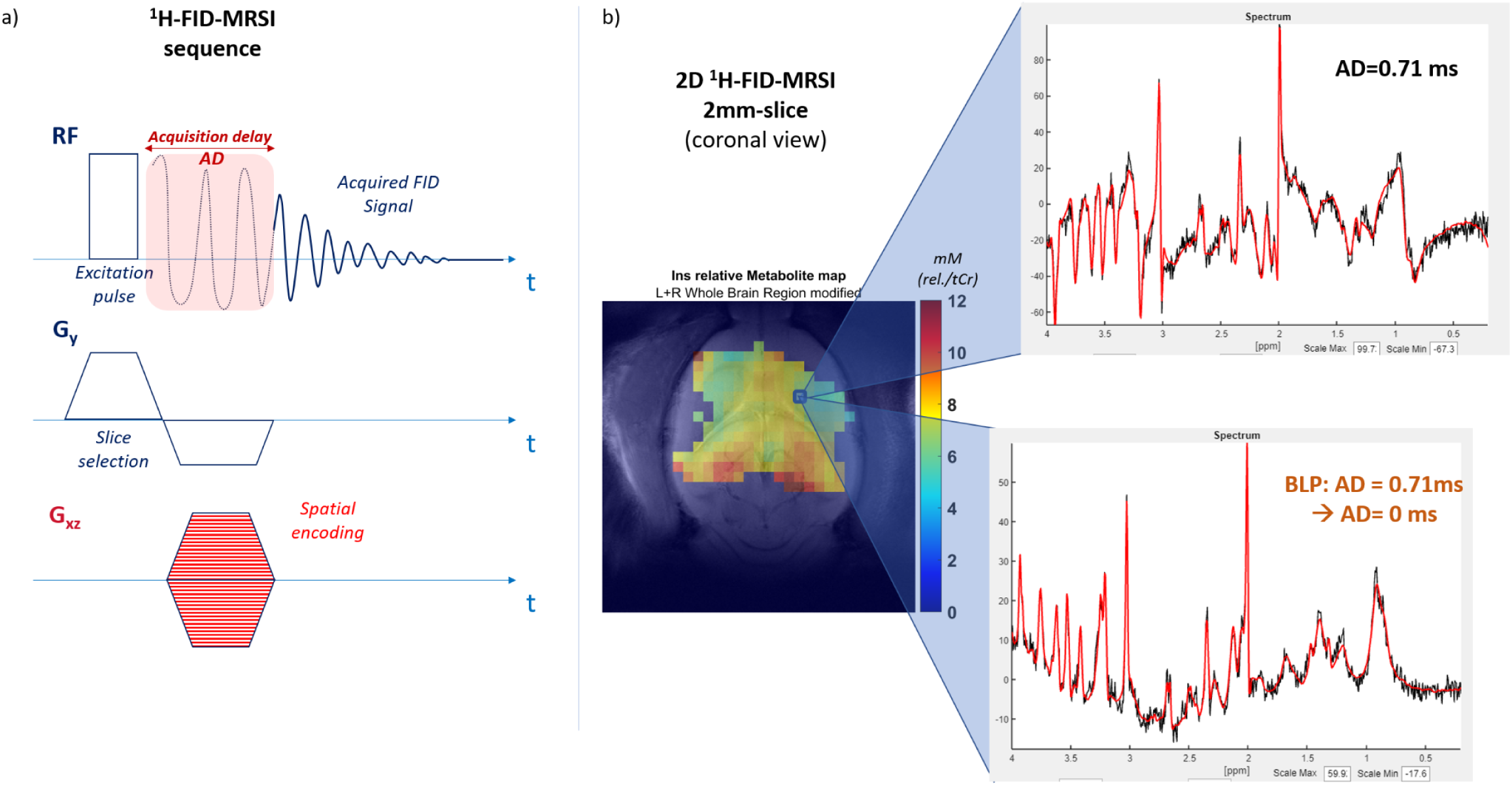
a) The FID-MRSI sequence scheme. The need for an acquisition delay (AD) between the excitation pulse and the FID acquisition window to spatially encode the signal causes a loss of signal. b) A representative acquired 2D ^1^H FID MRSI metabolic map (2 mm slice, 14.1T, Ins concentration). Each voxel concentration estimation arises from the relative ^1^H FID MRSI spectrum quantification: a representative in vivo spectrum acquired at AD = 0.71 ms is shown at the top, while at the bottom the reader can observe a representative backward-linear-predicted spectrum (AD = 0 ms).

^1^H-FID-MRSI studies have dealt with this feature in spectral fitting using three different approaches: a) by using a basis-set simulated with an AD matching the experimental configuration for all FIDs in the simulated metabolites [4] [10]; b) by simulating a basis-set at an AD = 0 ms setting, and excluding then the initial FIDs points to reach the experimental AD [5]; c) by predicting the missing initial points of the acquired FIDs using a backward linear prediction (BLP) autoregressive algorithm [6] [11]. The first two strategies allow the processing of the FID signal without relevant modifications, adapting the basis sets simulations only. However, despite LCModel [12] being one of the widely used spectral fitting methods, the impact of the AD in the metabolite estimation has never been investigated, especially with respect to the large and different first-order spectral dephasing induced by different AD values - and strong magnetic field strengths (B0). Indeed, LCModel was initially designed and validated for single-voxel MRS spectra [12], which do not require special handling of substantial first-order phase components.

The BLP methodology has proven useful in correcting spectral phase [11] [13], allowing a clearer visualisation of spectra. To date, no studies have systematically examined the impact of acquisition delay (AD) or the application of backward-linear-prediction (BLP) methods on metabolite quantification using LCModel in FID-MRSI.

In this work, we aim to address these gaps and provide new insights into the effects of AD and BLP in the context of FID-MRSI analysis. First, we present *in vivo* ^1^H-FID-MRSI data in the rat hippocampus, acquired with varying ADs to investigate whether AD introduces a bias in metabolite estimation. In parallel, we perform MC simulations of analogous ^1^H-FID-MRSI acquisitions for an extended analysis of quantification precision and biases.

We further investigate the impact of the BLP approach on metabolite quantification. MC simulations are employed to evaluate how the number of back-predicted FID points and the associated AD range influence estimation accuracy.For completeness, we also apply the BLP methodology to *in vivo* data, back-predicting FID points up to AD = 0 ms, and compare these results with the original acquisitions to evaluate the practical impact of BLP on metabolite concentration estimates.

## 2. Methods

### 2.1 ^1^H FID MRSI - Impact of varying ADs on metabolite estimates

#### 2.1.1 In vivo data

All experiments were approved by the Committee on Animal Experimentation for the Canton de Vaud, Switzerland (VD 3022.1, VD 3892). 2D ^1^H FID MRSI *in vivo* data were acquired in a shielded 14.1T scanner (Bruker/Magnex) on N = 3 Wistar rats (285 ± 25 g). The rats were anesthetized by 1.5%-2.5% isoflurane and their body temperature was kept at 37.5 ± 1.0 °C via a warm water circulation system. Both temperature and respiration were monitored during the acquisitions using a small-animal monitor system (SA Instruments, New York, NY, USA). MR data were acquired using a home-made transmit/receive quadrature surface coil (two 18×16 mm^2^ oval loops covering a curved 18×27 mm^2^ surface with a 14 mm radius of curvature). The first stage of the protocol [4] consisted of acquiring T2-weighted TurboRARE images (20 slices, 256 × 256 matrix size, RAREfactor = 6, axial and coronal orientations) for subject positioning and anatomical references. Shimming was performed using Bruker MAPSHIM in a 10 × 10 × 2 mm^3^ slab localized within the 2D FID MRSI slice of interest.

2D ^1^H-FID-MRSI acquisitions in a 2 mm thickness coronal slice, positioned in the brain and covering the hippocampus, the striatum and part of the cortex (Fig. 1b) were acquired for each rat with 3 different acquisition delays: AD = 1.30 ms, 0.94 ms and 0.71 ms, to investigate the effect of various ADs on spectral fitting and metabolites estimation (Fig. 2a). A Shinnar–Le Roux excitation pulse with a 8.4 kHz bandwidth and a 52° Ernst angle was employed. The FID acquisition was carried out with Cartesian k-space sampling, a 7.143 kHz acquisition bandwidth and 1024 spectral points, while the repetition time TR was set to 811 ms. The 2D MRSI 31 × 31 matrix size and 24 × 24 mm^2^ FOV resulted in a nominal voxel size of 0.77 × 0.77 × 2.00 mm³. No averaging was applied on the acquired FID signals and an in-plane 2D Hamming *k*-space filter was used (built in Bruker PV 360 v3.3). Water suppression was achieved withVAPOR [14] for metabolites signal acquisition, and 6 FOV saturation slices were used.

**Fig. 2:**
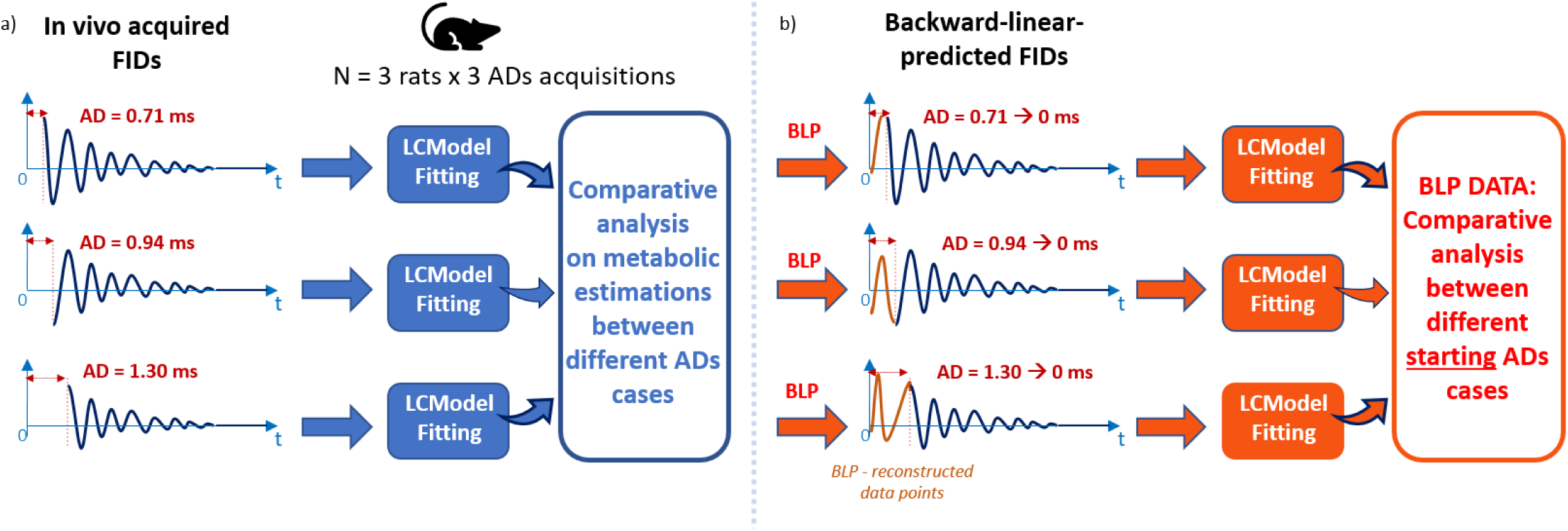
a) Illustrative diagram of the study steps on in vivo data to investigate the impact of varying ADs on metabolite estimates. On each rat (n=3), 3 ^1^H FID MRSI acquisitions with three different ADs (= 0.71 ms, 0.94 ms, 1.30 ms) were performed. After LCModel fitting, the metabolic concentration variations between the different ADs scenarios were investigated. b) The same ^1^H FID MRSI data acquired with different ADs underwent BLP to AD = 0 ms, LCModel fitting and an analogous comparative analysis as for original in vivo data.

#### 2.1.2 MC simulations

Monte Carlo simulations (n simulations = 1000) were performed to replicate 14.1T ^1^H-FID-MRSI rat brain spectra, under ideal conditions (i.e. no lipid contamination or water signal). These included ^1^H MR spectra of 18 metabolites (alanine-Ala, ascorbate-Asc, aspartate-Asp, creatine-Cr, phosphocreatine-PCr, γ-Aminobutyric acid-GABA, glutamine-Gln, glutamate-Glu, glutathione-GSH, inositol-Ins, lactate-Lac, N-acetylaspartate-NAA, taurine-Tau, glucose-Glc, N-acetylaspartylglutamate-NAAG, Phosphatidylethanolamine-PE, Glycerophosphocholine-GPC, posphatidylcholine-PCho) simulated with NMRScope-B/jMRUI [15] using an AD = 0 ms pulse-acquire sequence (FID time-domain points set to 1024 with a 7.143 kHz acquisition bandwidth as for *in vivo* data), considering literature chemical shifts and J-coupling inputs [16] [17], to which *in vivo* acquired macromolecules (double inversion recovery STEAM, TI1=2200ms, TI2=850ms, 10 x 10 x 2 mm^3^ voxel, metabolite residuals removed with AMARES [18]) were also added[19]. Each individual metabolite spectrum was then weighted to obtain realistic metabolic concentration values for the rat hippocampus based on existing data (Supplementary Table 1) [10] [20], and added together to represent a typical rat bain spectrum using an in-house built code (Matlab, MathWorks, Natick, MA, USA) [21] [22]. Gaussian noise was added in the spectral domain to generate 1000 realisations of realistic signal-to-noise ratio spectra (SNR = 18, defined as the ratio between the highest amplitude point of the real FID component and the noise amplitude σ, to match typical hippocampus MRSI values) [4] [10] [23].

All the n = 1000 FID signals were shortened of the first 5, 7 or 9 time-domain points (and zero-filled for the same length at their end, to keep a number of time-domain points of 1024) to obtain respectively the following AD scenarios: AD = 0.70 ms, 0.98 ms, 1.26 ms, approximations of in-lab used ^1^H-FID-MRSI AD values [4] [19]. Of note, the MC AD values were chosen to be multiple of the dwell time (0.14 ms), matching *in vivo* ADs as closely as possible whilekeeping the same experimental dwell time.

#### 2.1.3 Data processing and Statistical test

For the *in vivo* data, the MRS4Brain Toolbox [24] was employed to perform coronal brain segmentation, to process ^1^H-FID-MRSI data by removing the residual water signal via a Hankel singular value decomposition (HLSVD) algorithm and to apply semiautomatic quality control on voxel FIDs, fitted with LCModel. Such a quality control was based on LCModel-provided SNR (to be equal to or above 4), full width at half maximum (FWHM, to be lower than 125% of the average over the voxels) and Cramér–Rao lower bound (CRLB, to be lower than 40%). Furthermore, the toolbox allowed for anatomical segmentation of the hippocampus region.

For both MC simulations and *in vivo* data, the quantification via LCModel was carried out with the following quantification parameters: baseline stiffness *dkntmn* = 0.25, free zero-order phase adjustment, first-order phase set fixed to zero. The same jMRUI-simulated [15] basis sets were employed for MC simulations and *in vivo* data – one basis set per AD (0.71 ms, 0.94 ms and 1.30 ms), consisting of ^1^H MR spectra of 18 metabolites simulated using a pulse-acquire sequence with the same settings as the *in vivo* ¹H-FID-MRSI. For the MC simulations data, all the quantification outcomes atAD = 0.98 ms and AD = 1.26 ms were normalized as follows:, the metabolite concentration value was proportionally recalculated by setting the corresponding AD = 0.70 ms realisation outcome to concentration values of reference (Supplementary Table 1) [18] [19], to highlight the relative variations between different AD acquisitions. Analogously, for each *in vivo* MRSI hippocampus voxel, all the quantified metabolites concentration values were normalized to the AD = 0.71 ms case.

The whole ADs study focused on 6 metabolites of interest, whose concentration in brain ^1^H-FID-MRSI has proven to be estimated robustly [4]: Glu, Gln, Ins, Tau, tNAA, tCho, quantified with tCr as internal reference (set to 8 mM, (/tCr)). To detect possible quantification biases that are statistically significant between the different ADs outcomes, a Z-test was performed taking into account merely the quantified metabolic concentrations distributions for each AD, either over the quantified *in vivo* voxels orMC realisations. Furthermore, a t-test (n-2 degrees of freedom) was used in the analysis of simulation-by-simulation and voxel-by-voxel correlation. In particular, a linear regression was applied on the scatter plots comparing the concentration estimates over the different AD cases for each metabolite. A t-test was performed to determine whether the slope significantly deviated from 1, which represents the ideal consistency case (a unitary slope indicating perfect correspondence across voxels for in vivo data and across realisations for Monte Carlo simulations).

### 2.2 Impact of BLP on ^1^H-FID-MRSI metabolite estimates

#### 2.2.1 BLP method

The BLP method consisted of a home-made script (Matlab) based on the use of a specific Matlab function (*fillgaps* [25]). This function allows the back-reconstruction of a signal based on predictive autoregressive modelling, selecting automatically the model order that minimizes the Akaike information criterion [26]. The method is applied on transformed image-space level FIDs to back predict the first missing time-domain (complex) points, back-extrapolating them up to AD = 0 ms as displayed in Fig. 3 (step 2).

**Fig. 3:**
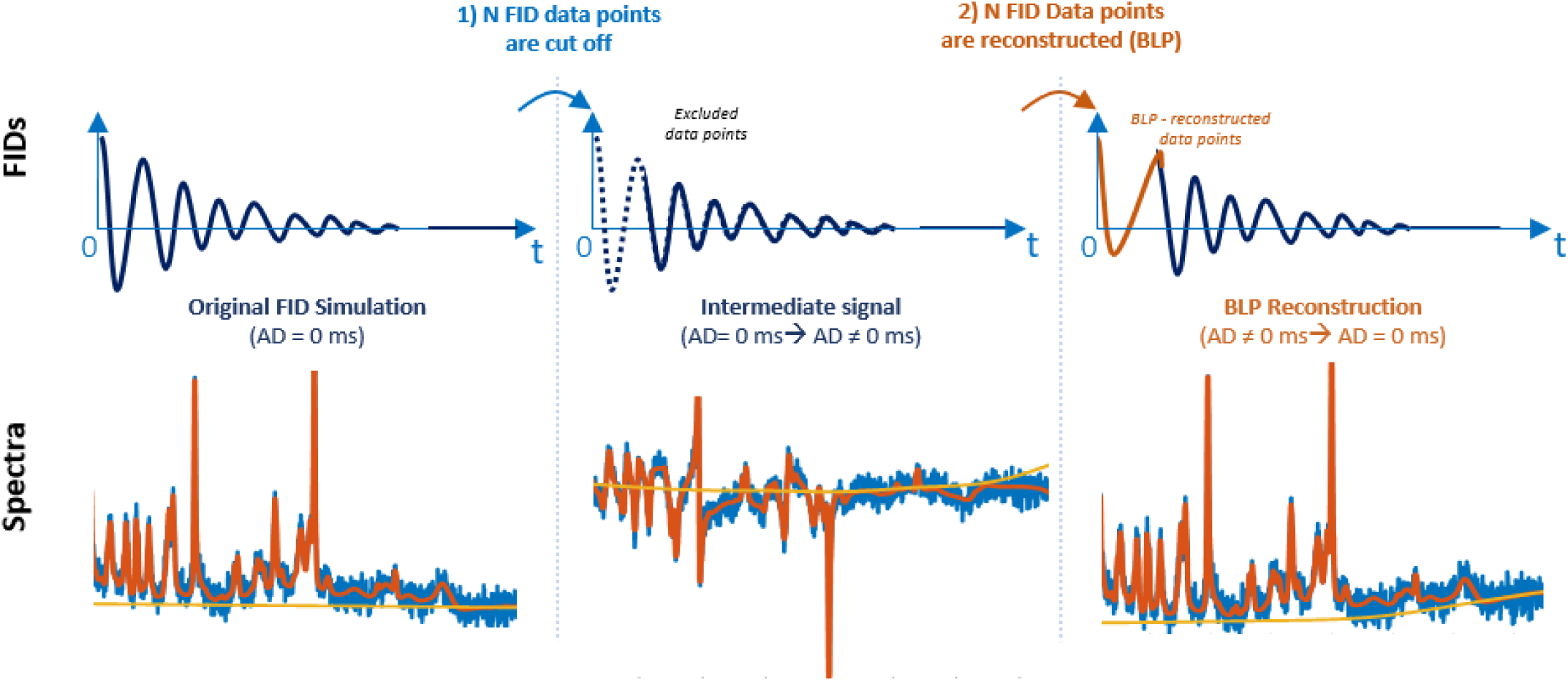
The cut-and-recover method via BLP tested in the consistency study introduced in paragraph 2.2.2. Applied on AD = 0 ms MC simulations, the first step consists of cutting off the first N FID data points to reach a certain AD ≠ 0 ms (e.g. in the figure a resulting spectrum at AD = 1.3 ms is displayed); the second step consists of the BLP algorithm application, that allows the back-prediction in the complex-domain of the first N missing FID points to reduce the AD up to 0. Such a step is applied also on in vivo ^1^H FID MRSI data acquired with AD = 1.30 ms, AD = 0.94 ms and AD = 0.71 ms, as illustrated in paragraph 2.2.3.

#### 2.2.2 BLP on MC simulations

Analogous ^1^H-FID-MRSI MC simulations (AD = 0 ms) to the ones described in paragraph 2.1.2 were performed to test the consistency of the BLP methodology. The simulations were carried out by setting an SNR = 12 (n simulations = 1000), to include a broader range of realistic MRSI SNR values *in vivo*. Different ADs scenarios were obtained by cutting off the first 2, 4, 7, 11, 15, 20, 30, and 40 FID time-domain points of each realisation, corresponding to AD values of 0.28, 0.56, 0.98, 1.5, 2.1, 2.8, 4.2, and 5.6 ms, respectively. The BLP method was applied in all cases to back-reconstruct the AD = 0 ms FIDs, in order to evaluate the consistency of this FID reconstruction method across increasing acquisition ADs. The obtained signals were fitted using LCModel with a basis set containing ^1^H MR metabolic spectral profiles, simulated with an AD = 0 ms pulse-acquire sequence in NMRScope-B/jMRUI [15]. A Z-test on the results of the different cut-and-recovered points cases was applied, by taking as reference the LCModel quantification outcome for a ground truth spectrum (i.e. 1 analogous realisation, AD = 0 ms, SNR = 500). The goal of such an analysis was to identify a potential maximum AD value for the application of BLP, (and the related number of back-predicted FID points) beyond which the reconstruction may fail for specific metabolites.

#### 2.2.3 BLP on in vivo data

The BLP method was further applied on the *in vivo* data introduced in paragraph 2.1.1 (n = 3 rats *x* 3 ^1^H-FID-MRSI acquisitions with AD = 1.30 ms, 0.94 ms and 0.71 ms, Fig. 2a-b) acquired on the same animals in the same conditions. The goal was to establish a common spectral quantification framework at a virtual delay point AD = 0 ms, at which the characteristic 1-st order phase evolution is eliminated. We back-predicted the missing FID time-domain points on the three ^1^H-FID-MRSI acquisitions (AD = 0.71 ms, 0.94 ms and 1.3 ms) up to AD = 0 ms, estimating respectively 5, 7 and 9 FID points. The BLP was performed on data preprocessed by an HLSVD algorithm (Matlab) for water residuals removal. LCModel fitting involved free zero-order phase management and first-order phase fixed to zero. A flexible baseline (*dkntmn* = 0.15) was used to account for possible baseline distortions previously mentioned in literature [13]. The data acquired at different ADs and quantified after back-prediction at AD = 0 ms were compared. To identify the best BLP quantification setup, 4 different modes were tested: 1) *full fitting range / MM,* fitting range set to 0.2 – 4.0 ppm with basis-set including measured MM; 2) *full fitting range / no MM,* fitting range set to 0.2 – 4.0 ppm with basis-set not including MM, 3) *partial fitting range / MM,* fitting range set to 1.8 – 4.0 ppm with basis-set including measured MM, 4) *partial fitting range / no MM,* fitting range set to 1.8 – 4.0 ppm with basis-set not including MM. The statistical analysis focused on the 6 metabolites of interest Glu, Gln, Ins, Tau, tNAA and tCho, on typical ^1^H-FID-MRSI segmented hippocampus data [4].

## 3. Results

### 3.1 ^1^H-FID-MRSI - Impact of varying ADs on metabolite estimates

#### 3.1.1 In vivo data

Fig. 4 (red bars) shows the mean quantification results for the *in vivo* analysis for the different ADs and metabolites, calculated over all the n = 113 hippocampus voxels under exam – aggregated from the N = 3 subjects of the study. The Z-test underlined the presence of scenario-by-scenario quantification biases. Statistically relevant discrepancies for all the AD transition cases were detected for the metabolites tCho (Fig. 4a, bottom), tNAA, Ins, Tau (Fig. 4b), while for Glu and Gln, mismatches were detected between the AD = 0.71 ms vs AD = 0.94 ms quantifications and the AD = 0.94 ms vs AD = 1.30 ms quantifications, respectively (Fig. 4a, top). The relative hippocampus quantification variations are shown in Table 1, and were particularly high ( > 10 %) for the AD = 0.71 ms vs AD = 0.94 ms comparison for tCho, tNAA and Tau (Table 1 on the right), and for Ins and tCho when comparing AD = 0.94 ms vs AD = 1.30 ms quantifications. A statistically significant discrepancy between the ideal case (i.e. same concentration estimates between the two ADs under exam, represented by a line with unitary slope) and the detected linear regression for the concentration estimates was observed for the following metabolites: tCho, Tau, tNAA in the AD = 0.71 ms vs AD = 0.94 ms comparison (Fig. 5a) and for Ins the AD = 0.94 ms vs AD = 1.30 ms case (Fig. 5c).

**Fig. 4:**
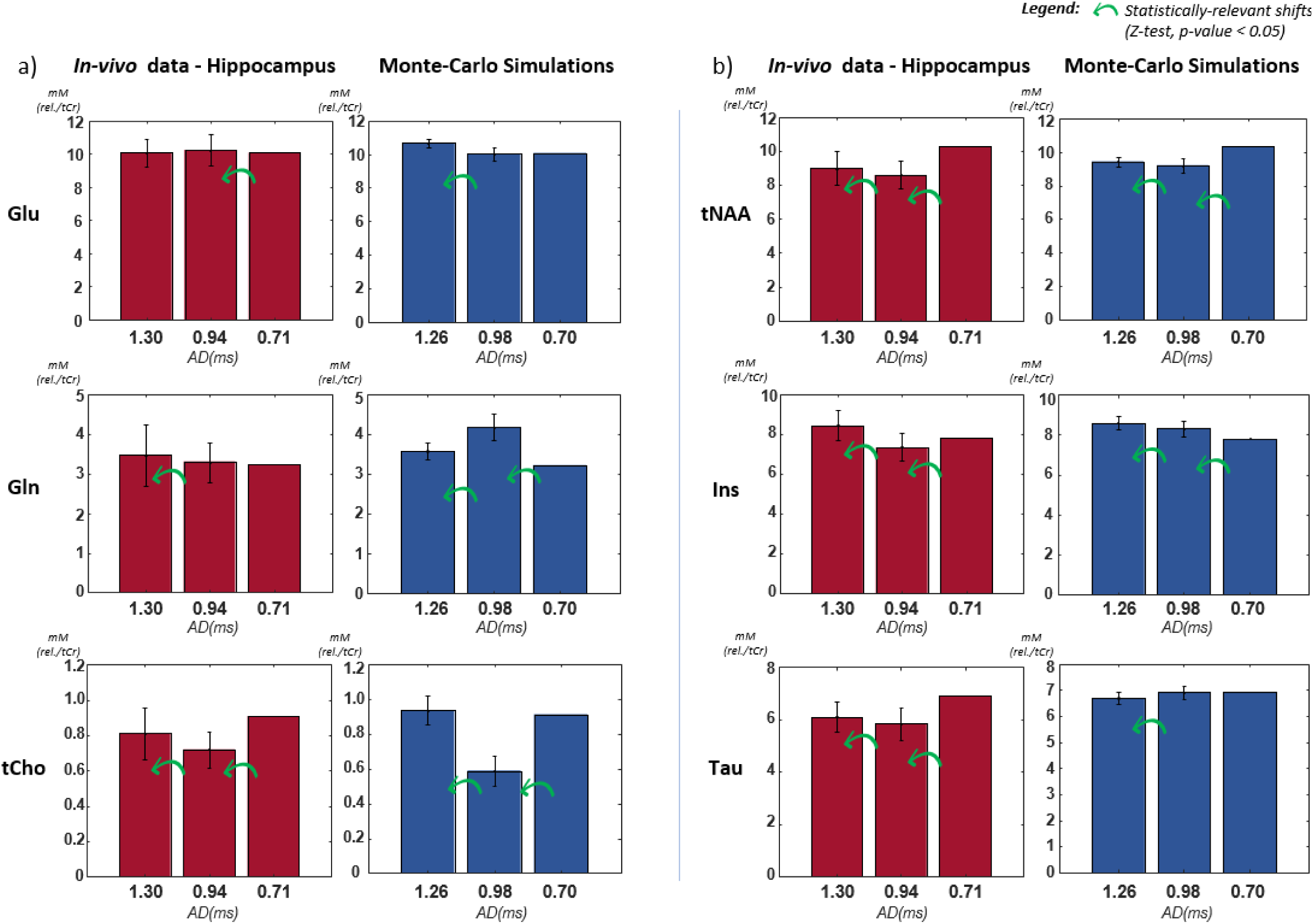
Mean quantification results at different ADs, including ^1^H FID MRSI in vivo data (red bar plots, n = 113 hippocampus voxels extracted from 3 rats) and MC simulations (blue bar plots, n = 1000 realisations) for the different examined AD scenarios. The metabolites of interest are Glu, Gln, tCho (a) and tNAA, Ins, Tau (b). All the bar plots show the calculated mean and standard deviation. Each in vivo voxel metabolite concentration estimate is normalized to its corresponding outcome at the shortest acquired AD (0.71 ms). The same procedure is applied for each MC realisation. The green arrows show statistically significant shifts of the concentration estimates for different ADs acquisitions and quantifications, with an AD-adjusted basis set (Z-test, p-value <0.05).

**Fig. 5:**
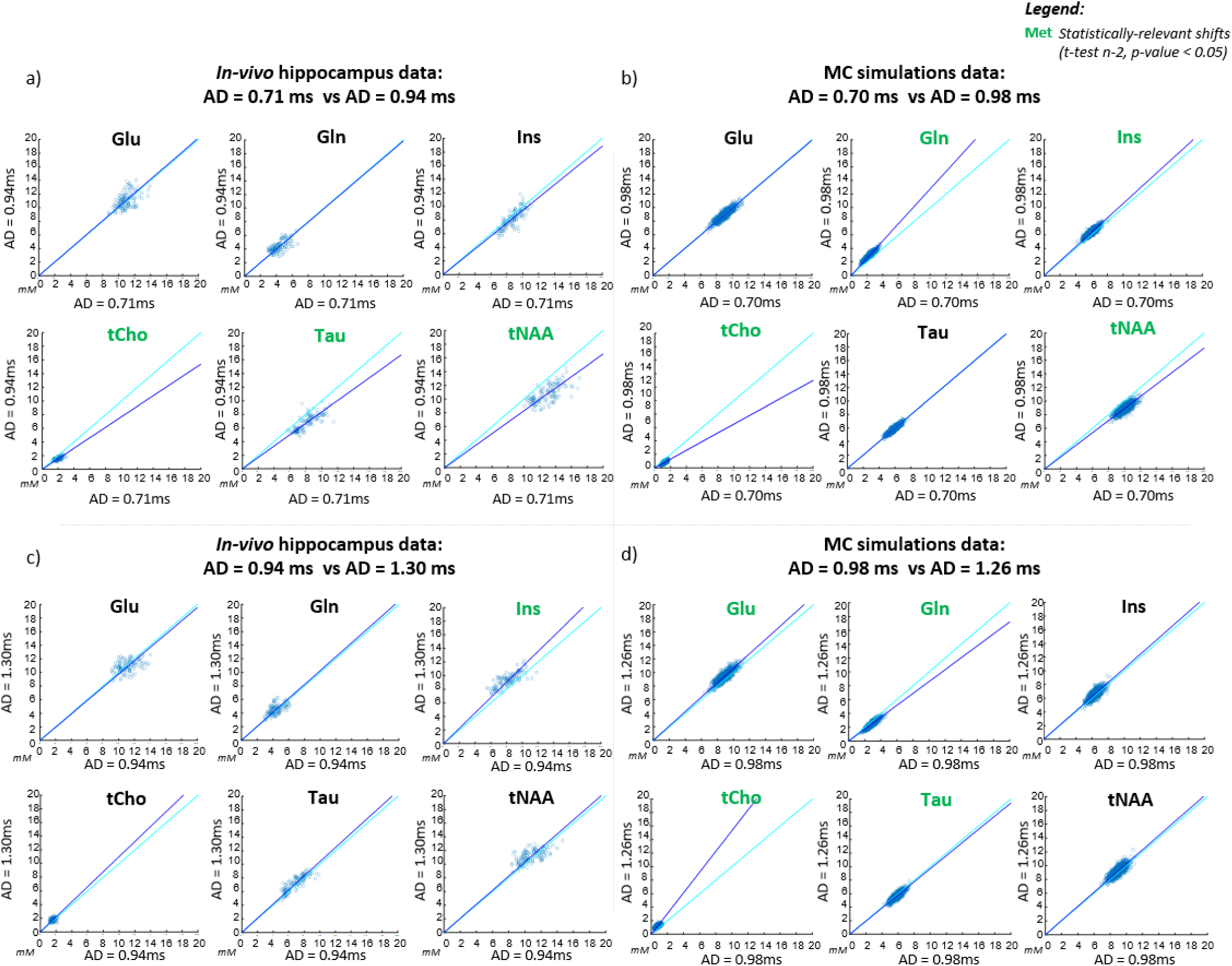
AD study correlation plots for Glu, Gln, Ins, tCho, Tau and tNAA. The displayed scatter values refer to: n = 113 hippocampus voxels the in vivo data in (a) and (c); n = 1000 realisations for the MC simulations in (b) and (d). The metabolites written in green showed a statistically significant shift (t-test with n-2 degrees of freedom, p-value < 0.05) with respect to the ideal case – i.e. absence of quantification differences between ADs, represented by an unitary slope line.

**Table 1:**
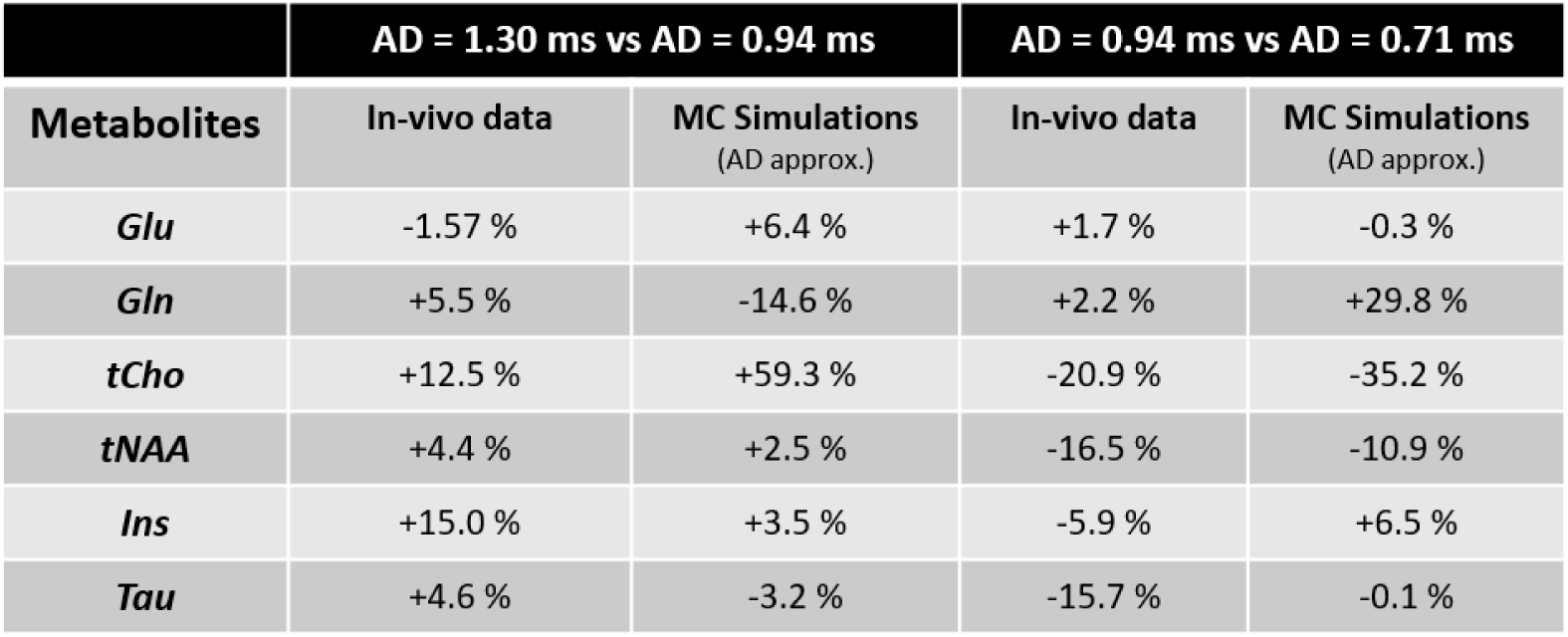
Mean metabolites quantification variations between two ADs acquisitions, with an AD-adjusted basis set, for in vivo data and MC simulations. For the latter, the displayed outcomes refer to scenarios whose AD values are the closest to the corresponding in vivo data parameters (e.g. the MC simulations AD = 1.26 ms value as an approximation of AD = 1.30 ms), set as a multiple of the dwell time for further BLP application.

|  | AD = 1.30 ms vs AD = 0.94 ms |  | AD = 0.94 ms vs AD = 0.71 ms |  |
| --- | --- | --- | --- | --- |
| Metabolites | In-vivo data | MC Simulations<br>(AD approx.) | In-vivo data | MC Simulations<br>(AD approx.) |
| *Glu* | -1.57 % | +6.4 % | +1.7 % | -0.3 % |
| *Gln* | +5.5 % | -14.6 % | +2.2 % | +29.8 % |
| *tCho* | +12.5 % | +59.3 % | -20.9 % | -35.2 % |
| *tNAA* | +4.4 % | +2.5 % | -16.5 % | -10.9 % |
| *Ins* | +15.0 % | +3.5 % | -5.9 % | +6.5 % |
| *Tau* | +4.6 % | -3.2 % | -15.7 % | -0.1 % |

#### 3.1.2 MC simulations

Fig. 4 (blue bars) shows the mean concentration values obtained fromthe MC simulations for each tested AD for Glu, Gln, tCho (Fig. 4a) and tNAA, Ins, Tau (Fig. 4b). Each simulation value is normalized to the corresponding lowest AD outcome (0.70 ms) - as described in paragraph 2.1.3. The analysis detected metabolite-specific estimation variations between the different AD-scenarios, and also between the AD ≠ 0 ms cases and the original AD = 0 ms simulation (Fig. 6). The corresponding relative discrepancies, displayed in Table 1, and were particularly high for tCho and Gln for both AD scenarios comparisons (AD = 0.98 ms vs AD= 1.26 ms and AD = 0.70 ms vs AD = 0.98 ms), with values higher than 10 %. Statistically significant mismatches (blue bars in Fig. 4, Z-test, green arrows: p-value <0.05,) were related to all the AD quantification comparisons for Gln, tCho, tNAA, Ins, while for Glu and Tau, they are detected between AD = 0.98 ms and AD = 0.70 ms only. Fig. 5b and 5d show the correlation plots obtained for all the 1000 simulations quantifications for AD = 0.70 ms vs AD = 0.98 ms and AD = 0.98 ms vs AD = 1.26 ms comparisons, respectively. An analogous t-test to the one applied on *in vivo* data was applied on MC simulations, underlining the presence of divergences in concentration estimates for the metabolites: Gln, Ins, tCho and tNAA in the case AD = 0.70 vs AD = 0.98 ms (Fig. 5b); Glu, Gln, tCho and Tau in the case AD = 0.98 ms vs AD = 1.26 ms (Fig. 5d).

**Fig. 6:**
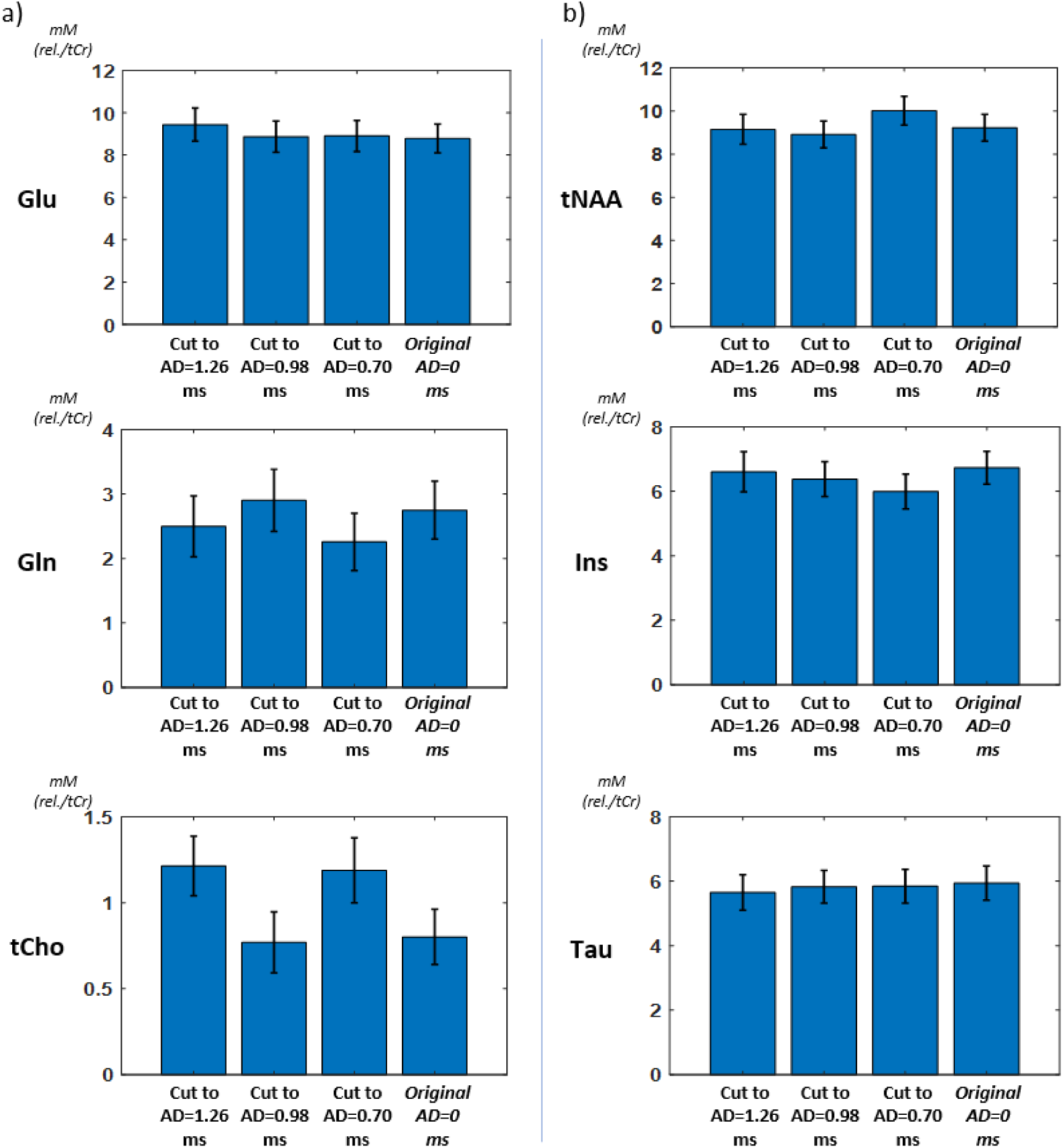
Mean quantification results at different ADs on MC simulations ( n = 1000 realisations) for the different examined AD scenarios and the original AD = 0 ms realisations. The metabolites of interest are Glu, Gln, tCho (a) and tNAA, Ins, Tau (b)., All the bar plots show the calculated mean and standard deviation. It is observable that the estimation results of the original MC simulations (AD = 0 ms) can vary when cutting FID points up to the tested ADs for most of the metabolites, particularly for Gln, tCho (a) and tNAA, Ins (b).

### 3.2 Impact of BLP on ^1^H FID MRSI metabolite estimates

#### 3.2.1 BLP on MC simulations

Fig. 7 shows metabolic quantification results of the cut-and-recover BLP method on original AD = 0 ms MC simulations of ^1^H FID MRSI signals. For all the metabolites, the mean concentration estimation retrieved on the 1000 realisations is displayed for all the cut-and-recovered FID points scenarios, as well as its standard deviation. The Z-test results showed statistical consistency over the whole examined AD range (0 - 5.6 ms) for the metabolites Tau and tCho (Fig. 7, bottom).

**Fig. 7:**
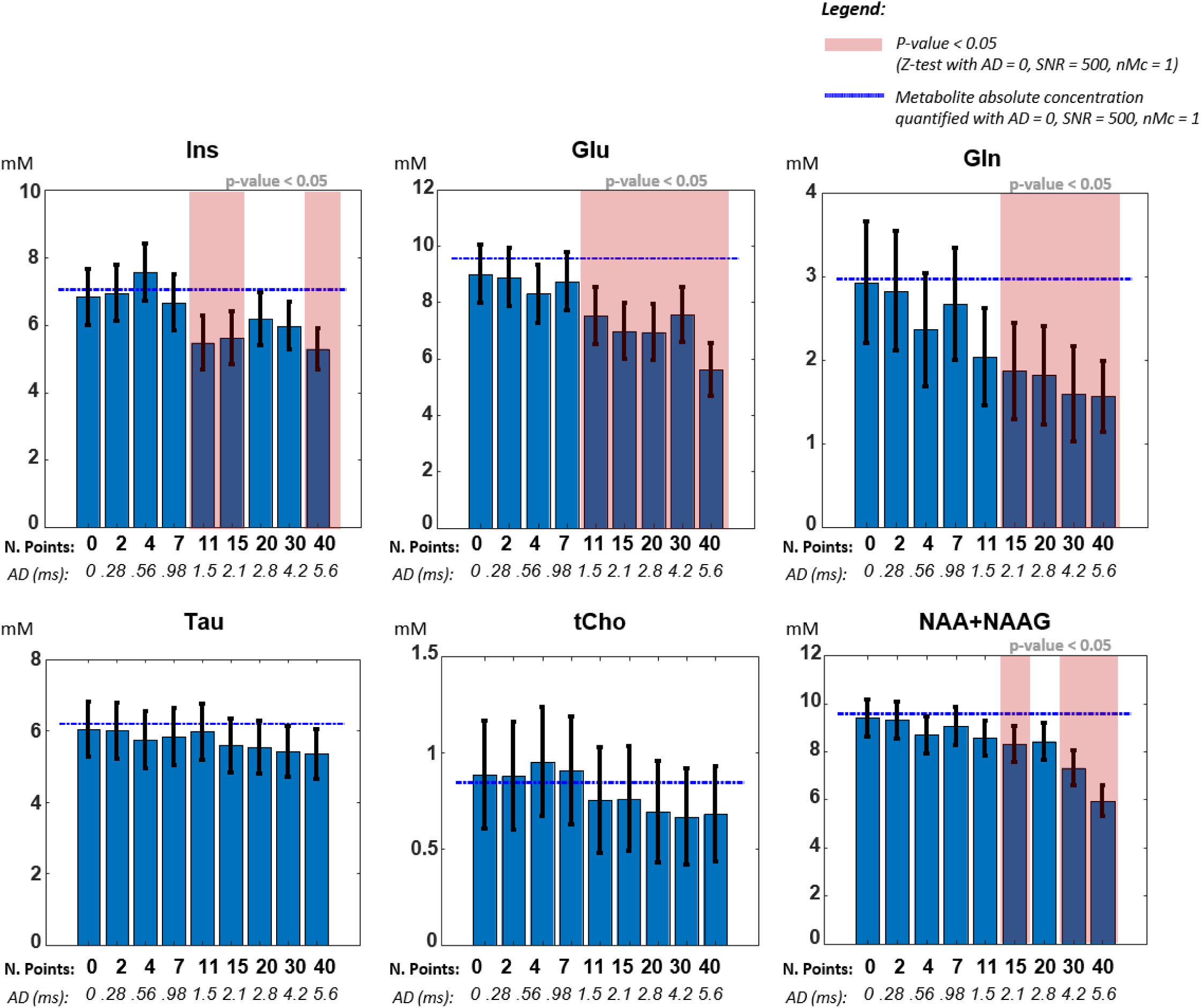
MC simulation results of the effect of BLP FID points recovery to a reference data with AD = 0 ms on metabolites quantification. The bar plots show mean and standard deviation of the LCModel quantification over 1000 MC simulations for increasing numbers of cut-and-recover points for the metabolites Ins, Glu. Gln. Tau. tCho. tNAA.

For the other tested metabolites, the consistency range covered AD = 0 - 0.98 ms for Ins and Glu (0-7 back-predicted FID points), AD = 0 - 1.50 ms for Gln and tNAA (0-11 back-predicted FID points). The ground-truth scenario taken as a reference for the Z-test is an SNR = 500 / AD = 0 ms MC realisation, whose quantification outcome is represented in Fig. 7 by a horizontal dashed blue line.

#### 3.2.2 BLP on in vivo data

The Table 2 lists all the four processing settings tested in the quantification of Backward-Linear-Predicted FID MRSI data at AD = 0 ms and their corresponding mean relative discrepancy when starting from AD = 0.94 ms vs AD = 0.71 ms acquisitions, for all the considered metabolites, in comparison with the original non-BLP data discrepancy (first row). The results show an overall average decrease of such quantification discrepancies when switching from a *full fitting range* (0.2 ppm to 4.0 ppm) to a *partial fitting range* (1.8 ppm to 4.0 ppm) settings. Such a trend is particularly visible for Glu, Tau, tNAA and tCho, while for Ins and Gln either a stable situation or an opposite light trend are observable. Overall, all the tested BLP settings (Table 2, last four rows) provided a mean discrepancy over Glu, Gln, Ins, Tau, tNAA and tCho between AD = 0.94 ms and AD = 0.71 ms acquisitions smaller than the non-BLP quantification scenario (Table 2, first row, average discrepancy of 10.5 %).

**Table 2:** BLP to AD = 0 ms application on in vivo data – relative discrepancy between BLP from AD = 0.94 ms and from AD = 0.71 ms hippocampus quantification values, in comparison with original in vivo data AD-related discrepancy. The tested settings included full fitting range (0.2 ppm to 4.0 ppm) and partial fitting range (1.8 ppm to 4.0 ppm) case, as well as basis-set with and without MM signal component included. All the tested BLP settings lead to positive results in terms of limiting the relative discrepancy between the tested AD cases, with a mean discrepancy over Glu, Gln, Ins, Tau, tNAA and tCho smaller than 10 %, obtaining a significant reduction in comparison with the original relative quantification difference (first line) between AD = 0.94 ms and AD =0.71 ms acquisition

| ADs comparison | LCModel settings | Glu | Gln | Ins | Tau | tNAA | tCho | Mean |
| --- | --- | --- | --- | --- | --- | --- | --- | --- |
| Original in-vivo data<br>AD = 0.94 ms vs<br>AD = 0.71 ms | Full range - MM | 1.7 % | 2.2 % | 5.9 % | 15.7 % | 16.4 % | 20.9 % | **10.5 %** |
| **BLP to AD = 0 ms:**<br>applied on<br>AD = 0.94 ms<br>vs<br>applied on<br>AD = 0.71 ms | Full range - MM | 8.5 % | 0.7 % | 1.0 % | 11.0 % | 6.3 % | 16.8 % | **7.4 %** |
|  | Full range - no MM | 6.6 % | 5.9 % | 0.9 % | 4.1 % | 4.6 % | 12.3 % | **5.7 %** |
|  | Partial range - MM | 4.7 % | 3.3 % | 5.5 % | 6.3 % | 0.2 % | 12.5 % | **5.4 %** |
|  | Partial range – no MM | 5.5 % | 3.7 % | 2.5 % | 4.1 % | 1.3 % | 10.5 % | **4.6 %** |

The BLP scenarios with the lowest discrepancy values are characterised by a *partial fitting range* data processing setting at AD = 0 ms, with a mean relative mismatch outcome of 5 % over all the metabolites of interest and over both tested basis sets - with and without MMs (Table 2, last two lines). For the *partial fitting range / no MM* LCModel setting, Fig. 8 shows the mean concentration estimates for all the investigated ADs data, BLP reconstructed and quantified at AD = 0 ms. While for the two lowest ADs (AD = 0.94 ms and AD = 0.71 ms) a level of divergence < 5% is detected for Gln, Ins, Tau, tNAA resulting in the already mentioned 4.6 % of mean relative mismatch, for AD = 1.30 ms acquisitions, quantification divergence is detected, characterised by a mean relative discrepancy of 15.7% over all the metabolites when compared to the AD = 0.94 ms acquisition.

**Fig. 8:**
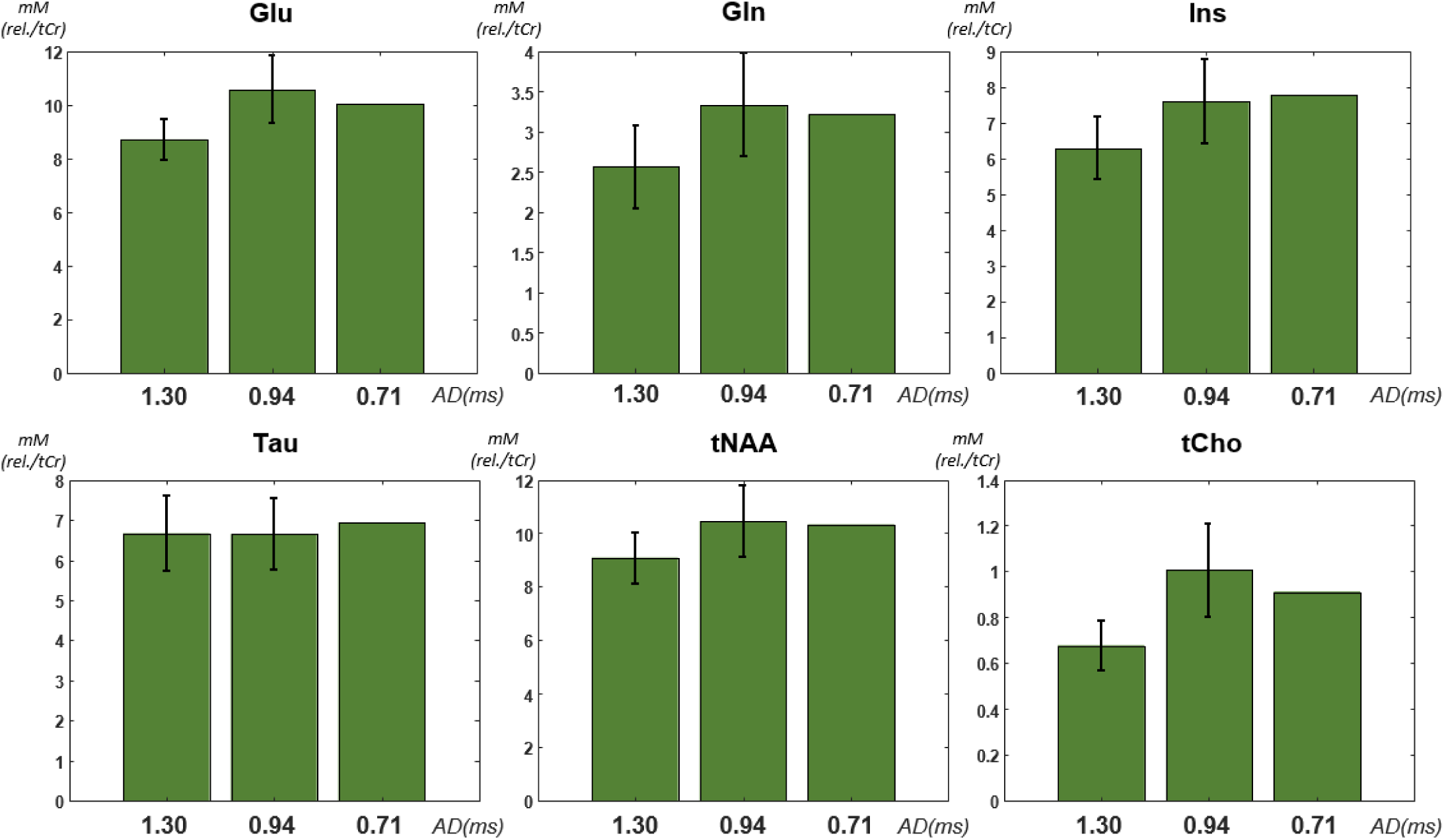
BLP validation study quantification results for the metabolites of interest. The bar plots show the calculated mean and standard deviation (n = 113 hippocampus voxels) for the different metabolites estimation for all the AD scenarios after the application of BLP up to AD = 0 ms. Each in vivo voxel metabolite estimate is normalized to its corresponding outcome at the lowest starting AD value scenario (0.71 ms), to highlight AD scenario-by-scenario variations at the voxel level. A significant convergence level of estimation outcomes for the two shorter ADs cases (AD = 0.94 ms and AD = 0.71 ms), while for the AD = 1.30 ms data a significant quantification divergence is observable for all the tested metabolites with the exception for Tau.

Fig. 9a illustrates the divergences obtained on representative metabolic maps from FID MRSI acquisitions and quantifications performed at different ADs on the same subject in the same scanning session. Ins, Tau, and tCho (Fig. 9a) show clear quantification divergences, as also highlighted by the relative mismatch maps. The application of the BLP methodology to back-predict FIDs up to AD = 0 ms (Fig. 9b) reduced the quantification divergences for Tau, NAA and tCho in most of the central area of the investigated brain slice.

**Fig. 9:**
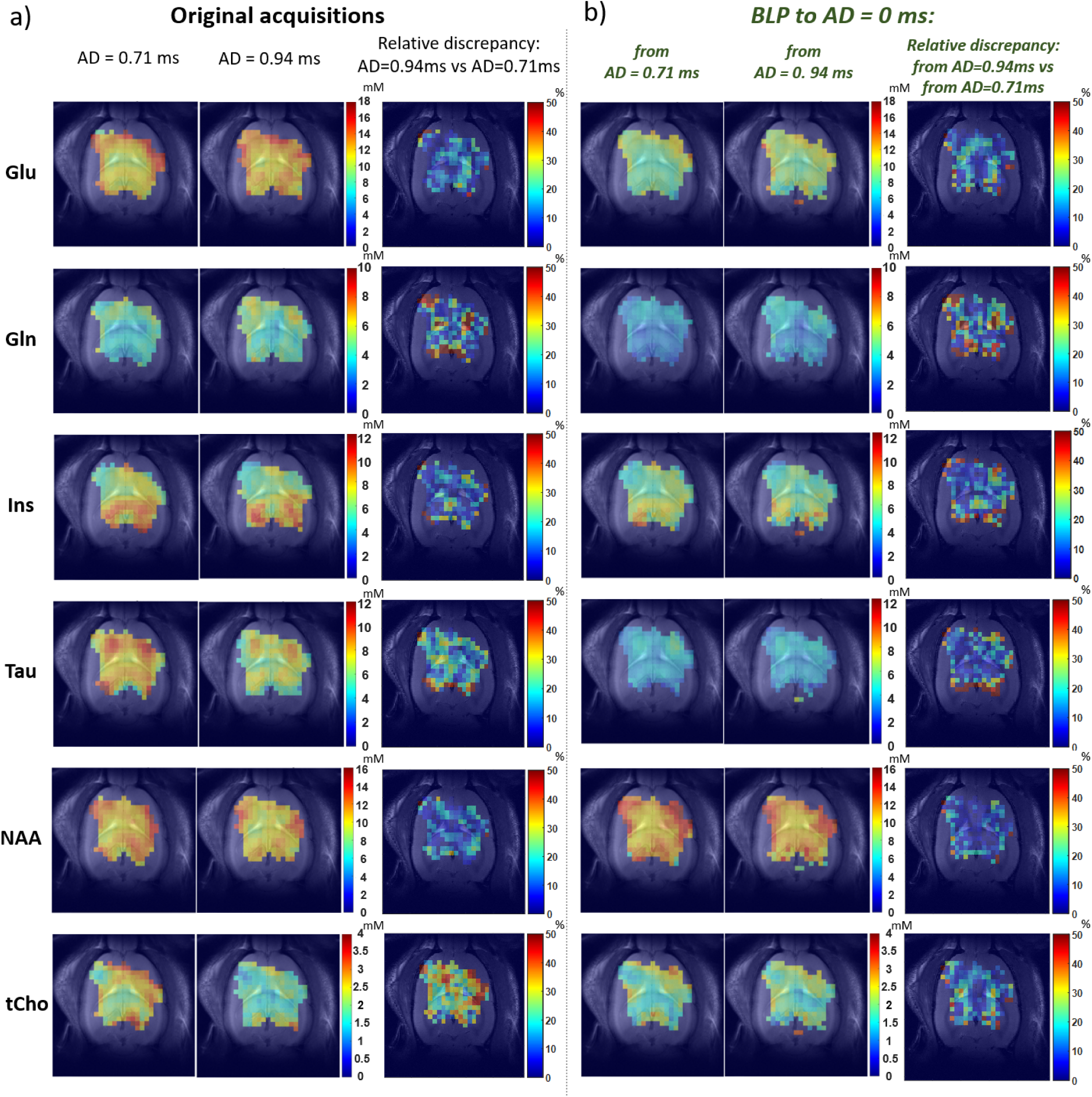
BLP application on in vivo data - Representative metabolic maps (mM rel./tCr). Divergent outcomes for FID MRSI acquisitions performed at AD = 0.71 ms and AD = 0.94 ms (a) converged when applying BLP to AD = 0 ms (b). This was particularly noticeable for Ins, Tau, tCho and NAA, as highlighted by the relative discrepancy maps for the original acquisitions (a) and BLP data (b).

## 4. Discussion

This work aimed to assess how the AD–induced first-order dephasing impacts on metabolite quantification in ¹H FID-MRSI, and to evaluate the potential of BLP to mitigate such biases by reconstructing spectra at a AD = 0 ms framework.

Both *in vivo* data and Monte-Carlo simulations demonstrated systematic AD-dependent quantification variations, with significant discrepancies for several metabolites despite the use of AD-specific basis sets.

BLP reconstruction showed high quantification consistency for a number of recovered FID points, enabling reliable estimation up to ∼AD = 0.98 ms for all the metabolites. *In vivo*, BLP substantially reduced inter-AD quantification mismatches—particularly for Tau, tNAA and tCho—when combined with a partial fitting range, lowering the mean discrepancy over all the metabolites from 10.5% to ∼5%.

### 4.1 ^1^H FID MRSI - Impact of varying ADs on metabolite estimates

#### 4.1.1 MC simulations

Fig. 4 (blue bars) and Fig. 6 illustrate that spectral quantification of truncated FIDs (and corresponding first-order dephased spectra) using an AD-specific metabolites basis set can lead to systematic quantification variations. For instance, for Glu (Fig. 4a, blue bars) and Tau (Fig. 4b, blue bars), while the transition from AD = 0.70 ms to AD = 0.98 ms did not imply relevant variations, the quantification at AD = 1.26 ms introduced a statistically significant quantification bias compared to AD = 0.98 ms. The analysis over other metabolites showed that different AD acquisitions and quantifications led to increased variations in concentration estimates, as underlined statistically by the applied Z-test (Fig. 4, blue bars). This was reflected into the relative discrepancies calculation shown in the Table 1, where significant mismatches (> ±5%) were observed in the MC study for Glu, Gln, tCho in the AD = 0.98 ms vs 1.26 ms comparison, and for Gln, tCho, tNAA, Ins in the AD = 0.70 ms vs 0.98 ms comparison. Furthermore, analysing the correlation plots between quantifications at different ADs and the related t-test outcome (Fig. 5), a statistical divergence between the ideal 1-to-1 correspondence and the detected linear regression was observed for most of the examined metabolites: Gln, Ins, tCho and tNAA in the case AD = 0.70 vs AD = 0.98 ms (Fig. 5b); Glu, Gln, tCho and Tau in the case AD = 0.98 ms vs AD = 1.26 ms (Fig. 5d).

Considering that a variation in the AD causes a first-order phase variation in the FID MRSI spectra, we can hypothesize that LCModel (originally designed for single-voxel spectroscopy) could struggle in handling such a strong first-order dephasing in the quantification process even though the metabolite basis set is matching the MC conditions. This hypothesis is also supported by the fact that these quantification biases also appeared on MC simulations performed in ideal conditions, i.e. without including lipids contamination nor water residuals, excluding their respective impact in the detected metabolites estimation biases. Of note, at 14.1T and for a typical AD of 1 ms, the first-order phase wrap is of 1080° over the 5ppm spectral range of interest.

#### 4.1.2 In vivo data

The *in vivo* analysis revealed clear quantification divergences for several metabolites, as shown in the correlation plots of Fig. 5a and 5c, with significant discrepancies for tCho, Tau and tNAA in the AD = 0.71 ms vs AD = 0.94 ms comparison (Fig. 5a) and for Ins in the AD = 0.94 ms vs AD = 1.30 ms case (Fig. 5c) (t-test, n–2, p-value < 0.05). These *in vivo* biases, statistically relevant for most metabolites examined – such as Glu and Gln (Fig. 4a) and Ins and Tau (Fig. 4b), Z-test – likely arise from non-idealities intrinsic to *in vivo* spectra, including residual water and lipids, which are absent in MC simulations and may themselves vary with AD. Such effects can modulate the LCModel handling of first-order phase contributions, thereby influencing the quantification trends. When comparing these *in vivo* outcomes with MC simulations (Table 1), analogous behaviours were observed for tCho and tNAA, consistent with Fig. 4 (red bars), whereas other metabolites showed different patterns with respect to simulated conditions. Overall, the *in vivo* findings support the discrepancies highlighted in the MC study (paragraph 4.1.1) and reinforce the hypothesis of a potential limitation in LCModel when processing strongly first-order dephased FID-MRSI spectra.

### 4.2 Impact of BLP on 1H FID MRSI metabolite estimates

#### 4.2.1 BLP on MC simulations

Fig. 7 shows that the BLP reconstruction had a measurable impact on metabolites quantification, with a strong consistency observed for Tau and tCho over the whole examined AD range (0 - 5.6 ms). For the other metabolites, such a consistency was reduced to a range of AD = 0 - 0.98 ms for Ins and Glu (0-7 back-predicted FID points) and to AD = 0 - 1.50 ms for Gln and tNAA (0-11 back-predicted FID points). This underlined the potential of the BLP method as a tool in overcoming LCModel quantification biases at AD ≠ 0, for a maximum backwards reconstructed FID of 7 BLP FID points, corresponding to AD = 0.98 ms. It is important to note that this limit was derived under specific conditions of magnetic field strength (14.1 T) and signal-to-noise ratio (SNR = 12), which closely align with typical ^1^H FID-MRSI acquisitions performed in our group. Furthermore, such a limit underlines that AD values as 0.71 ms or 0.94 ms (as performed in our group) in combination with BLP methodology, offer a promising alternative for FID-MRSI spectral quantification with minimal bias.

Back-calculated spectra at AD = 0 ms also demonstrated a high degree of consistency in spectral recovery, with visible biases only appearing when more than 20 BLP points were recovered (AD = 2.8 ms, Fig. 10). The measured quantification divergence occurring as the number of recovered points increases may be linked to spectral baseline oscillations that could arise from imperfectly predicted initial FID points. This would correspond to the addition of an artificial fast decaying component in the FID, translated as a very broad component in the spectral domain. Indeed, a trend of a slightly rising baseline component was progressively observed (Fig. 10) for increasing BLP recovered points, which could potentially introduce quantification biases across various metabolite components. However, such minor variations in the baseline can be easily accommodated in the spectral quantification with a reasonable degree of flexibility of the baseline in the LCModel fitting, by setting the corresponding flexibility parameter *dkntmn* accordingly – as described above in paragraph 2.2.3.

**Fig. 10:**
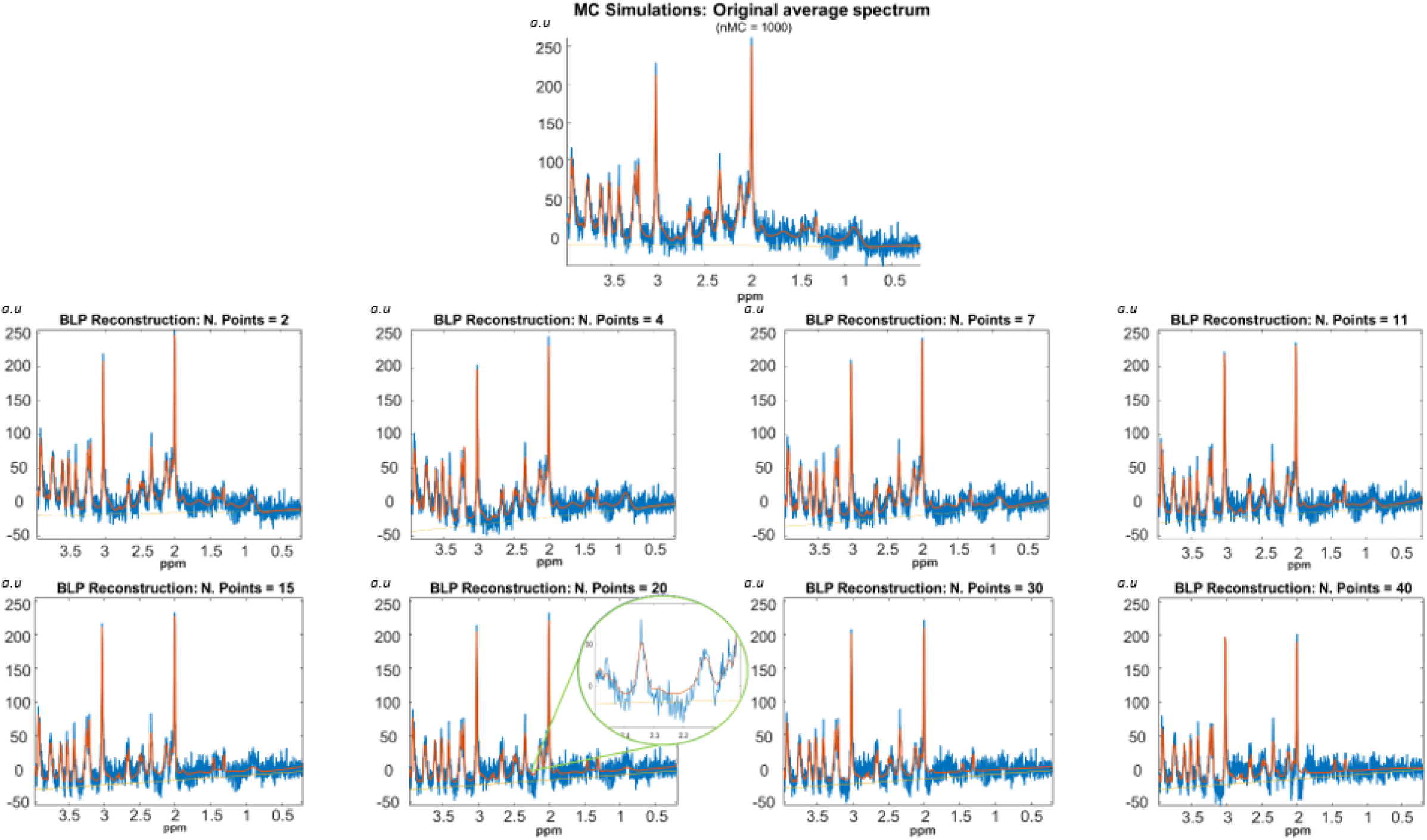
Examples of simulated spectra (blue) and their LC Model fitting (orange) obtained from the Monte-Carlo simulations. A reference spectrum is first simulated for AD = 0 ms and then cut for a certain number of points (2 to 40, thus from 0.28 ms to 5.6 ms), back-reconstructed with the BLP algorithm to AD = 0 ms and quantified with LC Model. As the number of cut points rises, an increasing mismatch between data and fitting function starts to be visible in the range of 20 – 40 points.

#### 4.2.2 BLP on in vivo data

Besides potential BLP-induced baseline variations, the impact of the MM profile in FID-MRSI quantification was investigated, by testing basis-sets with measured MM and without MM. A partial fitting range (4.0 ppm – 1.8 ppm [5] [9]) was also considered, to test for a possible better quantification robustness towards potential lipid contamination and reduced impact of potential BLP-induced baseline variations in the metabolite characterization. As displayed in Table 2, the relative discrepancy between quantifications of BLP-reconstructed data to AD = 0 ms from AD = 0.94 ms and AD = 0.71 ms in the rat hippocampus was limited to a beneficially low range, with a mean discrepancy over Glu, Gln, Ins, Tau, tNAA and tCho smaller than 10 %. The best consistency was obtained for the *partial fitting range* results (Table 2, last 2 rows), where the mean relative discrepancy calculated over the examined brain metabolites in *MM* and *no MM* cases was 5.4% and 4.6%, respectively.

Comparing such an outcome with the original FID-MRSI quantifications discrepancy (AD = 0.94 ms vs AD = 0.71 ms, Table 2, first row) underlines the general positive impact of such a BLP processing strategy. Indeed, considering the setting “*partial fitting range/ no MM”* (Table 2, last line), besides a slight deterioration in Glu and Gln estimation (going from an average discrepancy of 2.0% to 4.6 %) massive improvement in the characterization of Tau (from ∼ 16% to 4%), tNAA (from ∼ 16% to 1%) and tCho (from ∼ 21% to 11%) were obtained. Ins relative discrepancy was reduced as well (from 5.9% to 2.5%). Additionally, the BLP approach in this case reduced the average relative quantification divergence to 4.6 % instead of the original 10.5 % over the investigated brain metabolites.

Overall, analysis of the results obtained under the different processing settings at AD = 0 ms (Table 2) indicates that using a narrower fitting range of 4.0–0.2 ppm improves LCModel quantification of BLP FID-MRSI data. Interestingly, the BLP quantifications with partial fitting range (4.0 ppm – 1.8 ppm, and without MM inclusion) provided the best agreement between the acquisitions done at AD = 0.94 ms and AD = 0.71 ms in identical conditions. This can be related to the fact that the dominant MM resonance lies outside this range (0.9 ppm peak), making MM identification challenging in the fitting process and therefore affecting the quantification variability. Furthermore, all tested BLP settings (Table 2, last four rows) effectively reduced the relative discrepancy between the evaluated AD cases.

Applying the *partial fitting range / no MM scenario* data quantification setting, Fig. 8 underlined the good level of convergence of quantification outcomes for the two shorter ADs scenarios (AD = 0.94 ms and AD = 0.71 ms), while for the AD = 1.30 ms data, a significant quantification divergence (15.7% with the AD = 0.94 ms acquisition) was observed. This is coherent with the MC study discussed in the paragraph 4.2.1, where a BLP consistency range value was identified up to an AD value of 0.98 ms (and 7 FID points), thus excluding the AD = 1.30 ms in the presented cases. Another interesting agreement with the MC study can be found for the Tau quantification results, which do not diverge significantly in the AD = 1.30 ms acquired data case (Fig. 8, bottom), in agreement with the good level of consistency measured for Tau for the whole investigated AD range (0 – 5.6 ms) in the MC study (Fig. 7, bottom). It is also worth noting that an AD value choice is not necessarily driven by the dwell time, inducing thus the BLP application to leave some first order dephasing residual, corresponding to a residual delay smaller than half the dwell time.

The potential of the BLP method application and spectral quantification at a virtual AD = 0 ms delay on *in vivo* data is also illustrated by the metabolic maps obtained over the whole rat brain. The representative metabolite concentration maps in Fig. 9 illustrate marked visual differences between FID-MRSI acquisitions analyzed at different ADs (Fig. 9a). These discrepancies were eliminated when applying BLP at AD = 0 ms with quantification using a common basis set (Fig. 9b). This effect was particularly evident for Ins, Tau, tCho, and NAA.

Overall, these *in vivo* results highlight the capability of BLP method in overcoming initial LCModel quantification discrepancies caused by the intrinsic AD in an FID MRSI sequence and the related first-order spectral dephasing. It is worth underlining that this holds also for *partial fitting range / MM* processing setting: MM profiles have been shown to play a relevant role in the brain metabolites characterization with SVS [19] and their correct inclusion are also expected to improve the FID-MRSI quantification. Measured MM have typically shown better quantification outcomes for SVS and are expected to perform as well for FID-MRSI, although their absolute characterization is still a challenge.

Our results pave the way for the BLP application on *in vivo* ^1^H FID-MRSI studies, enabling the use of fully phased spectra within a unified analysis framework and a single basis set at AD = 0 ms for acquisitions with varying delays. This standardization facilitates easier cross-study comparisons. Moreover, the generation of phased spectra greatly enhances the visual interpretability of FID-MRSI data, supporting clearer identification of spectral peaks, amplitude variations, and potential contaminations or distortions. Finally, the resulting MRSI spectra are restored to a phasing state consistent with SVS quantifications, where validated quantification methods and analysis strategies are known to perform reliably.

To conclude, all evaluated BLP quantification strategies (whether using a full or reduced ppm range and including or excluding MM components in the basis set) consistently improved spectral quantification across data acquired at different ADs, with stable performance over a wide range of voxels. It should be noted that this analysis was conducted under 14.1 T magnetic field conditions, and variations may occur under different field strengths due to changes in first-order phase behavior.

Future work will focus on a more detailed evaluation of incorporating MM components into the basis set for BLP data quantification, given their established role in SVS-based metabolite characterization [27]. Additionally, further investigation into applying BLP to water signals for water-based metabolite estimation is warranted. Extending BLP to X-nuclei, particularly in the context of direct deuterium MRSI characterized by short T₂ values, could be especially promising. In such cases, the impact of similar ADs and the associated loss of initial FID points on spectral information is expected to be more pronounced, where BLP may offer significant potential for recovering FID signal integrity.

## 5. Conclusion

This study demonstrates that varying AD in ^1^H FID-MRSI significantly influencesinfluence metabolite concentration estimates when spectra were quantified using LCModel with AD-adapted basis sets. Both *in vivo* data and Monte Carlo simulations revealed systematic quantification biases, underscoring the need to carefully consider AD effects in FID-MRSI fitting.

The BLP methodology showed a good level of consistency in MC simulations up to the AD value of 0.98 ms in the FID points back-predicting at 14.T and realistic SNR conditions. Importantly, BLP enabled the harmonization of the divergent results obtained from AD = 0.94 ms and AD = 0.71 ms data by quantifying them within a common basis set framework at AD = 0 ms. This capability facilitates cross-study comparisons and supports efforts toward MRSI standardization. Additionally, visualizing spectra and quantification under both zero- and first-order fully phased conditions proved highly beneficial for improving spectral interpretation and data quality assessment.

## Acknowledgements

We acknowledge Dr. Jessie Mosso for providing coding resources to perform the Monte-Carlo simulations used in this study. We acknowledge the CIBM Center for Biomedical Imaging for providing expertise and resources to conduct this study. Financial support was provided by the Swiss National Science Foundation (Projects No. 201218 and 207935).

## Data Availability

The detailed datasets generated during and/or analysed during the current study are available from the corresponding author on reasonable request.

## Compliance with Ethical Standards

This study was funded by the Swiss National Science Foundation (Grants No. 201218 and 207935). All the authors declare they have no conflict of interest. All animal experiments were conducted according to federal and local ethical guidelines, and the protocols were approved by the local Committee on Animal Experimentation for the Canton de Vaud, Switzerland (VD 3022.1, VD 3892).

---

**SUPPLEMENTARY Table 1:** Concentration values used for the generation of MC simulations and for all the normalization procedures. These values were collected from in vivo measurements in the rat brain[20].

| Ala | Asc | Asp | Cr | PCr | GABA | Gln | Glu | GSH | Ins | Lac | NAA | Tau | Glc | NAAG | PE | GPC | PCho |
| --- | --- | --- | --- | --- | --- | --- | --- | --- | --- | --- | --- | --- | --- | --- | --- | --- | --- |
| 0.64 | 2.49 | 1.74 | 4.26 | 4.99 | 1.28 | 3.22 | 10.05 | 0.89 | 7.80 | 1.33 | 9.52 | 6.93 | 1.00 | 0.80 | 2.49 | 0.50 | 0.41 |

